## Supplementary File 4 for "Discovering molecular features of intrinsically disordered regions by using evolution for contrastive learning"

**Classification Datasets**

To assess if representations were capturing functional aspects of IDRs, we established three sets of classification datasets, containing a total of 117 binary classification problems.

First, we obtained two binary classification datasets (previously published by Zarin *et al*. (Zarin *et al.*, 2020)) predicting if IDRs are mitochondrial targeting signals or contain Cdc28 phosphorylation sites, respectively. Labels in the mitochondrial targeting signal IDR dataset correspond to data from a proteome-wide screen (Vögtle et al., 2009): a positive label “1” is a mitochondrial targeting sequence identified in this screen, while all other IDRs are assigned the negative label “0”. In total, there are 186 positive and 5168 negative IDRs in this dataset. Labels in the Cdc28 phosphorylation site IDR dataset correspond to high-confidence Cdc28 phosphorylation sites previously curated from *in vitro* and *in vivo* studies (Nguyen, Co, Irie, & Li, 2000). A positive label “1” is an IDR that overlaps any of these curated sites, while all other IDRs are assigned “0”. In total, there are 256 positive and 5109 negative IDRs in this dataset.

Second, as mitochondrial targeting and Cdc28 phosphorylation alone are not fully representative of the range of functions carried out by IDRs, we assessed agreement with computationally predicted functions for yeast IDRs in a previous bioinformatics analysis (Zarin *et al.*, 2019). Using 82 literature-curated features, the authors defined 23 clusters enriched in functional annotations according to a clustering of these features. We converted these into 23 classification problems, where “1” indicates membership in the cluster, and “0” is assigned to all other IDRs. Although these labels are at the IDR level, they are biased towards functions discoverable with the features used in this previous analysis.

Third, as the previous classification problems predict only a limited subset of IDR functions, measured by specific experimental assays or resolvable with features known to be associated with IDR function, sought to test if our features were predictive of function in general, without prior association of these functions with IDRs. To do this, we measured if proteins with the same GO annotations were clustered together according to their IDR features. Exploratory analyses often propagate biological knowledge from an IDR or protein’s nearest neighbors in a feature space; this principle can be used to generate hypotheses for IDRs of unknown function (Zarin *et al.*, 2019), or to find more examples of rare protein phenotypes by looking at the neighbors of the few known instances (Lu *et al.*, 2019). In line with these types of analyses, we implemented nearest neighbor classifiers for 92 GO Slim annotation terms, representing all terms that had more than 50 proteins represented in the subset of proteins with IDRs in our benchmarks. Unlike our previous datasets, GO Slim annotations are only available at a protein level, not for individual IDR function: to handle this, we averaged the feature representations of all IDRs together for each protein. Proteins were assigned to the positive set if its nearest neighbor had that GO Slim annotation, and to the negative set otherwise. We then measured the fold enrichment of this positive set for the annotation relative to background.

To ensure a fair benchmark, we used only the intersection of sequences being capable of being represented by all feature representation methods in all three benchmarks. We built logistic regression classifiers (using 5-fold cross validation) and nearest-neighbor classifiers (leave-one-out validation). For our logistic regression classifiers, we report the balanced accuracy and the AUC. For our nearest neighbor classifiers, we report the balanced accuracy and the precision. For the cluster annotation datasets, we report average metrics over all 23 classification problems. For the GO Slim annotation datasets, there is no expectation of predictive performance, since the functions we attempt to classify may not actually be mediated by IDRs. We instead sought to assess if proteins carrying out the same function are more clumped in the representation space than random: we report the fold enrichment of the GO annotation in the positive set compared to background. We report the median of this metric over all 92 classification problems. Classifiers were implemented with the scikit-learn package (Pedregosa *et al.*, 2012).

**Baseline Methods**

Baseline feature representations from previous self-supervised learning methods were preprocessed and calculated using the best practices described in their respective original work. For the SeqVec baseline, we took the sum of the embeddings across LSTM layers and averaged this sum over amino acid positions (Heinzinger et al., 2019). For the TAPE and Unirep baselines, we used the average embedding across amino acid positions (Rao et al., 2019).

For our literature-curated features (Zarin et al., 2019), we directly obtained a csv of the raw features calculated using only the *Saccharomyces cerevisiae* sequence directly from the authors (this feature set differs from the features reported in the original manuscript, which are the average and variance of all species against a simulated expectation.)

We trained two models from scratch using the same yeast dataset we trained our reverse homology model on. First, we trained a SeqVec model from random initialization for 100 epochs on this data using the implementation provided by the authors (Heinzinger et al., 2019). Second, we trained a weakly-supervised classification model, where the model takes an input IDR sequence and predicts the homologue set the sequence is from, using an identical encoder to our $g_{2}$ encoder with a fully-connected layer outputting the class prediction at the end. This model was trained using a standard categorical cross-entropy loss for 1000 epochs with a learning rate of 1e-4. Finally, we report results from a randomly initialized, untrained neural network with an identical architecture as our reverse homology implementation. Since $g_{1}$ and $g_{2}$ are identical in architecture minus the averaging over sequences for $g_{1}$ (which we do not use for post-training encoding in the benchmarks and interpretations in this paper), we simply took representations from $g_{2}$ from our random model.

**Benchmark Results**

We report metrics for each of our three sets of classification datasets as Supplementary Table 1, 2, and 3, respectively. We focus on representations from the final convolutional layer of our target encoder in the manuscript, as this layer yields interpretable features compared to the fully-connected layers, and encodes single sequences compared to the source encoder. However, for these benchmarks, we show performance for representations learned by the final convolutional and fully connected layers of both our source and target encoders. For features from convolutional layers, we express them as max and average pooled features across the entire sequence, as is done in our architecture natively. We compared these representations to literature-curated features and to representations from other self-supervised protein models based on natural language processing (NLP) strategies. The former contains features collected from almost three decades of IDR studies (Zarin *et al.*, 2019), while the latter are from universal protein representation models (Alley *et al.*, 2019; Heinzinger *et al.*, 2019; Rao *et al.*, 2019) typically trained on four orders of magnitude more data than our model with an order of magnitude more parameters.

We compared representations from a randomly initialized untrained model, to confirm that performance is dependent on training with our self-supervised proxy task. We also trained a model on a weakly supervised version of our training task, where the homology family is directly supplied as a label instead of being used to pose a contrastive learning problem, to confirm that performance depends on the contrastive structure of our proxy task, not just on supplied labels for homology. Finally, we compared representations from one other self-supervised protein method (SeqVec (Heinzinger *et al.*, 2019)) trained from random initialization on the same yeast IDR data we used to train our reverse homology model.

Supplementary Table 1. Performance of representations on classifying mitochondrial targeting signal IDRs and IDRs containing Cdc28 phosphorylation sites, with nearest-neighbor and logistic regression classifiers. We report standard error.

|  | Mitochondrial Targeting | | | | Cdc28 Phosphorylation | | | |
| --- | --- | --- | --- | --- | --- | --- | --- | --- |
|  | Nearest Neighbor | | Log. Regression | | Nearest Neighbor | | Log. Regression | |
|  | Bal. Acc. | Precision | Bal. Acc. | AUC | Bal. Acc. | Precision | Bal. Acc. | AUC |
| Reverse Homology  (target, conv3) | 0.617  ± 0.007 | 0.243 ± 0.006 | 0.761 ± 0.006 | 0.806 ± 0.005 | 0.591 ± 0.007 | 0.165 ± 0.005 | 0.782 ± 0.006 | 0.863 ± 0.005 |
| Reverse Homology  (target, fc2) | 0.640 ± 0.007 | 0.340 ± 0.006 | 0.761 ± 0.006 | 0.813 ± 0.005 | 0.590 ± 0.007 | 0.197 ± 0.005 | 0.789 ± 0.006 | 0.872 ± 0.005 |
| Reverse Homology  (source, conv3) | 0.626 ± 0.007 | 0.297 ± 0.006 | 0.744 ± 0.006 | 0.800 ± 0.005 | 0.568 ± 0.007 | 0.138 ± 0.005 | 0.781 ± 0.006 | 0.861 ± 0.005 |
| Reverse Homology (source, fc2) | 0.614 ± 0.007 | 0.252 ± 0.006 | 0.754 ± 0.006 | 0.794 ± 0.005 | 0.576 ± 0.007 | 0.159  ± 0.005 | 0.788 ± 0.006 | 0.863 ± 0.005 |
| Literature-curated | 0.591 ± 0.007 | 0.204 ± 0.006 | 0.716 ± 0.006 | 0.797 ± 0.005 | 0.556 ± 0.007 | 0.228 ± 0.006 | 0.796 ± 0.005 | 0.885 ± 0.004 |
| Sequence Similarity | 0.510 ± 0.007 | 0.096 ± 0.004 |  |  | 0.540 ± 0.007 | 0.127 ± 0.005 |  |  |
| TAPE | 0.647 ± 0.007 | 0.323 ± 0.007 | 0.753 ± 0.006 | 0.819 ± 0.005 | 0.582 ± 0.007 | 0.173 ± 0.005 | 0.785 ± 0.006 | 0.869 ± 0.005 |
| SeqVec | 0.671 ± 0.006 | 0.379 ± 0.007 | 0.784 ± 0.006 | 0.865 ± 0.005 | 0.611 ± 0.007 | 0.214 ± 0.006 | 0.788 ± 0.006 | 0.878 ± 0.005 |
| UniRep | 0.664 ± 0.006 | 0.354 ± 0.007 | 0.805 ± 0.005 | 0.900 ± 0.004 | 0.613 ± 0.007 | 0.217 ± 0.006 | 0.801 ± 0.005 | 0.897 ± 0.004 |
| Random  Features | 0.540 ± 0.007 | 0.127 ± 0.005 | 0.710 ± 0.006 | 0.769 ± 0.006 | 0.554 ± 0.007 | 0.103 ± 0.004 | 0.747 ± 0.006 | 0.811 ± 0.005 |
| Weakly Supervised | 0.605 ± 0.007 | 0.244 ± 0.006 | 0.691 ± 0.006 | 0.739 ± 0.006 | 0.574 ± 0.007 | 0.131 ± 0.005 | 0.763 ± 0.006 | 0.845 ± 0.005 |
| SeqVec (Yeast IDRs) | 0.633 ± 0.007 | 0.272 ± 0.006 | 0.752 ± 0.006 | 0.825 ± 0.005 | 0.558 ± 0.007 | 0.114 ± 0.004 | 0.762 ± 0.006 | 0.831 ± 0.005 |

Supplementary Table 2. Average performance of representations in classifying 23 previous computationally annotated clusters by Zarin *et al*., with nearest-neighbor and logistic regression classifiers. We report standard error.

|  | Nearest Neighbor | | Logistic Regression | |
| --- | --- | --- | --- | --- |
|  | Balanced Accuracy | Precision | Balanced Accuracy | AUC |
| Reverse Homology  (target, conv3) | 0.611 ± 0.007 | 0.234 ± 0.006 | 0.756 ± 0.006 | 0.832 ± 0.006 |
| Reverse Homology  (target, fc2) | 0.617 ± 0.007 | 0.239 ± 0.006 | 0.786 ± 0.006 | 0.856 ± 0.005 |
| Reverse Homology  (source, conv3) | 0.582 ± 0.007 | 0.181 ± 0.006 | 0.754 ± 0.006 | 0.844 ± 0.005 |
| Reverse Homology (source, fc2) | 0.581 ± 0.007 | 0.165 ± 0.005 | 0.776 ± 0.006 | 0.853 ± 0.005 |
| Literature-curated | 0.583 ± 0.007 | 0.210 ± 0.006 | 0.778 ± 0.006 | 0.852 ± 0.005 |
| Sequence Similarity | 0.528 ± 0.007 | 0.124 ± 0.005 |  |  |
| TAPE | 0.612 ± 0.007 | 0.239 ± 0.006 | 0.767 ± 0.006 | 0.840 ± 0.005 |
| SeqVec | 0.607 ± 0.007 | 0.240 ± 0.006 | 0.793 ± 0.006 | 0.868 ± 0.005 |
| UniRep | 0.602 ± 0.007 | 0.234 ± 0.006 | 0.777 ± 0.006 | 0.861 ± 0.005 |
| Random  Features | 0.569 ± 0.007 | 0.160 ± 0.005 | 0.747 ± 0.006 | 0.814 ± 0.006 |
| Weakly Supervised | 0.595 ± 0.007 | 0.182 ± 0.006 | 0.706 ± 0.007 | 0.782 ± 0.006 |
| SeqVec (Yeast IDRs) | 0.610 ± 0.007 | 0.218 ± 0.006 | 0.788 ± 0.006 | 0.856 ± 0.005 |

Supplementary Table 3. Median fold enrichment of representations for GO Slim annotations in nearest neighbor classifiers, using 92 GO Slim annotations.

|  | Median Fold Enrichment |
| --- | --- |
| Reverse Homology (target, conv3) | 1.917 |
| Reverse Homology (target, fc2) | 2.659 |
| Reverse Homology (source, conv3) | 1.623 |
| Reverse Homology (source, fc2) | 2.062 |
| Literature-curated | 1.872 |
| TAPE | 2.550 |
| SeqVec | 2.502 |
| UniRep | 2.360 |
| Random  Features | 1.523 |
| Weakly Supervised | 1.839 |
| SeqVec (Yeast IDRs) | 1.792 |

We observe that features from the final fully connected layer of our target sequence encoder perform better than features from the convolutional layers, or from the source sequence encoder. This result is unsurprising in the context of how the model was trained. The source encoder is trained to produce representations that will be averaged across several species, so just using it to represent single sequences out-of-context as we did in these evaluations is not consistent with its training (in contrast, the target encoder is used to represent single sequences during training.) The output of the fully connected layers is used for the contrastive proxy task, so we expect that the representations from this layer are most discriminative. These results indicate that the choice of layer to use in downstream tasks will depend on whether the application is interpretation or classification/prediction of sequences. However, we note that the features from the target convolutional layer still outperform at least the randomly trained models and the literature-curated features at all tasks, so the trade-off may not be drastic.

Our feature representations perform better than an untrained random baseline at all tasks, confirming that this performance is dependent on training with our self-supervised proxy task. They also outperform features from a model trained on a weakly supervised version of our reverse homology task, where the homology family is directly supplied as a training label instead of being used to pose a contrastive learning problem, indicating that contrastive learning is important to the quality of the learned features. Finally, our feature representations also outperform literature-curated features (Zarin *et al.*, 2019), along with the other self-supervised methods, suggesting that features learned by these neural networks may encode more functional information than expert-curated features.

Overall, these results suggest that our model represents IDRs at a similar level of performance as state-of-the-art self-supervised protein representation methods, but with a more constrained, low-parameter architecture and substantially less training data.
