## Supplementary figures and images for "Discovering molecular features of intrinsically disordered regions by using evolution for contrastive learning"

### filter_0.png

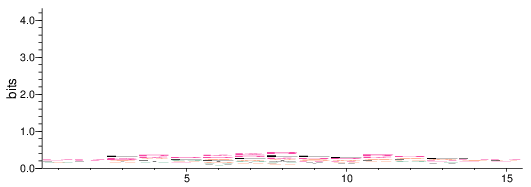

### filter_1.png

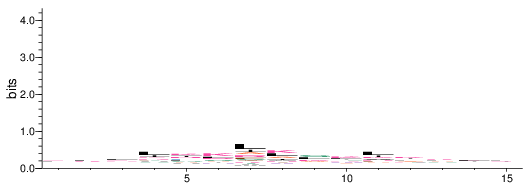

### filter_10.png

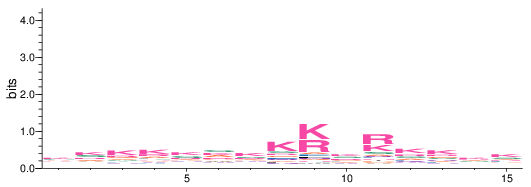

### filter_11.png

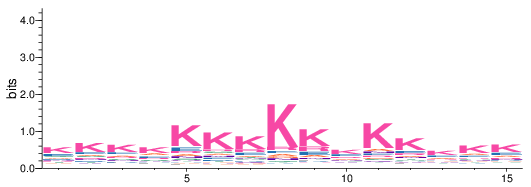

### filter_12.png

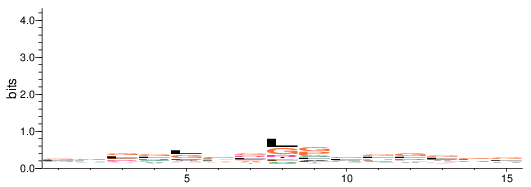

### filter_13.png

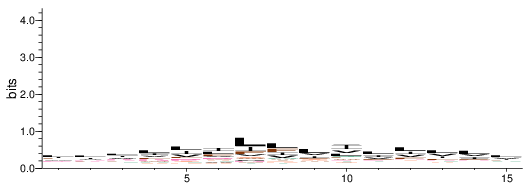

### filter_14.png

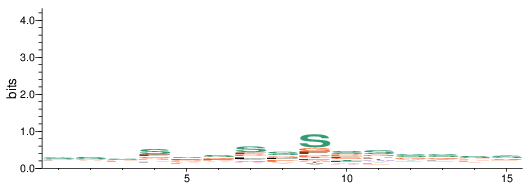

### filter_15.png

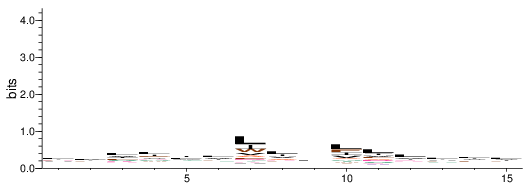

### filter_16.png

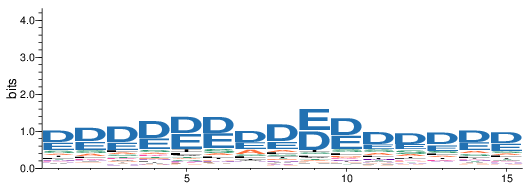

### filter_17.png

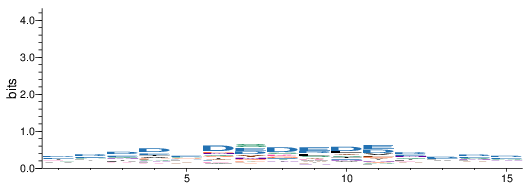

### filter_18.png

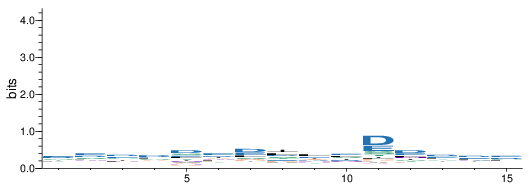

### filter_100.png

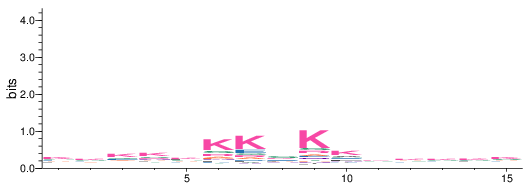

### filter_101.png

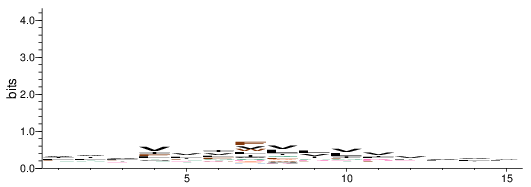

### filter_102.png

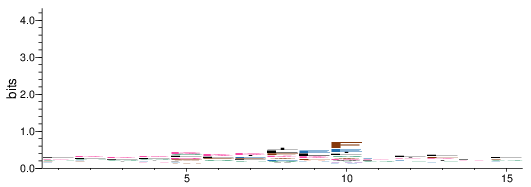

### filter_103.png

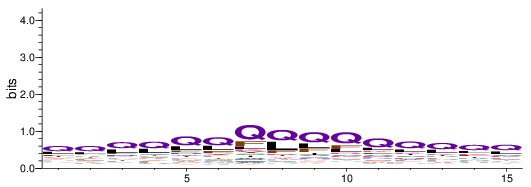

### filter_104.png

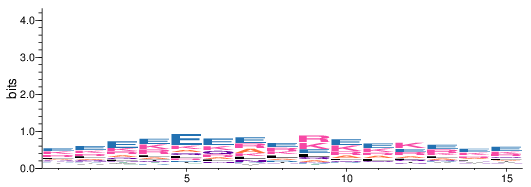

### filter_105.png

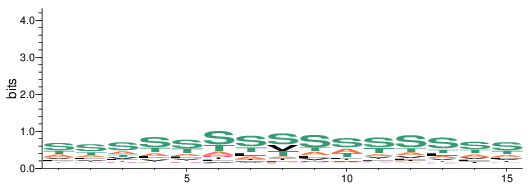

### filter_106.png

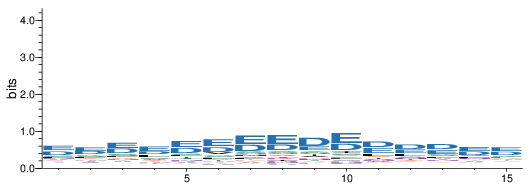

### filter_107.png

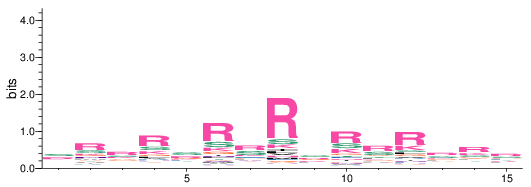

### filter_108.png

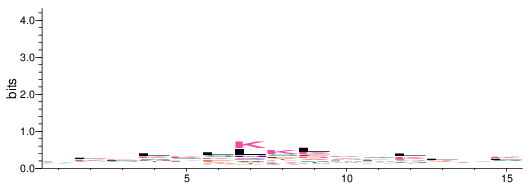

### filter_109.png

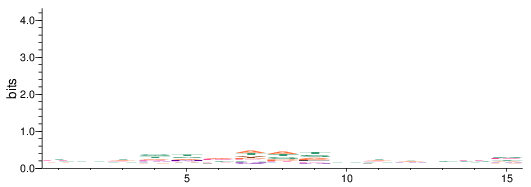

### filter_110.png

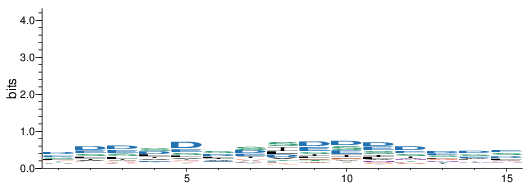

### filter_111.png

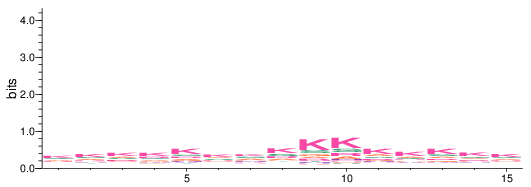

### filter_112.png

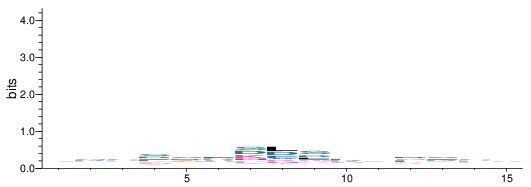

### filter_113.png

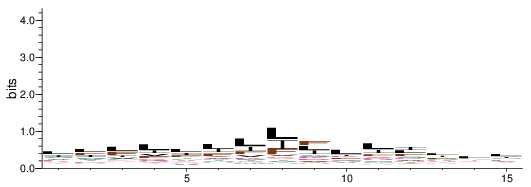

### filter_114.png

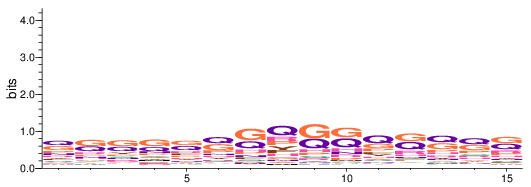

### filter_115.png

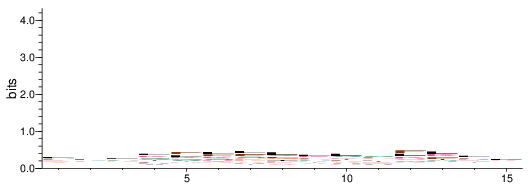

### filter_116.png

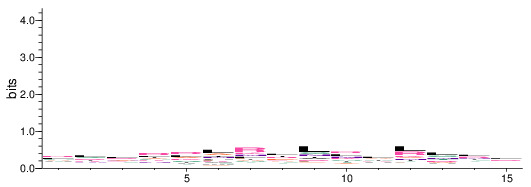

### filter_117.png

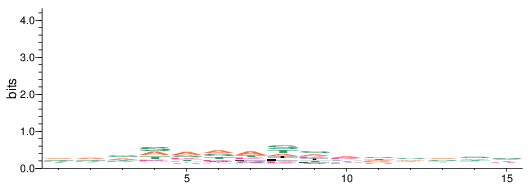

### filter_118.png

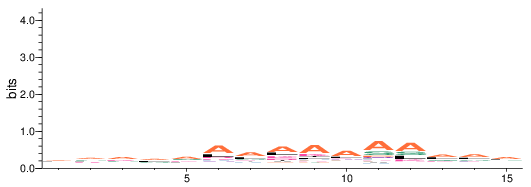

### filter_119.png

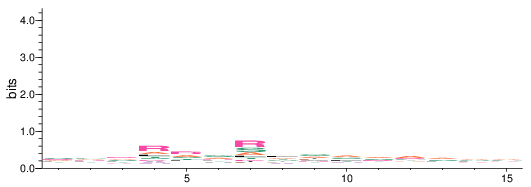

### filter_120.png

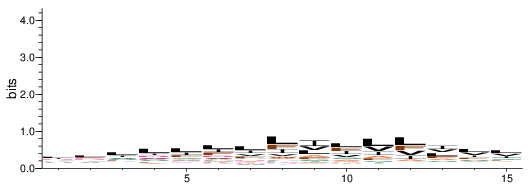

### filter_121.png

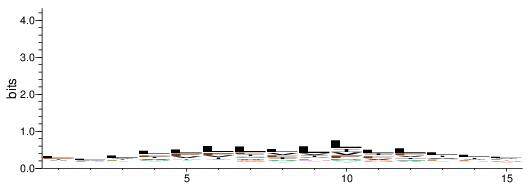

### filter_122.png

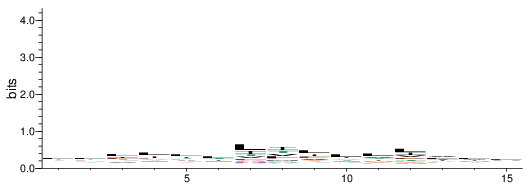

### filter_123.png

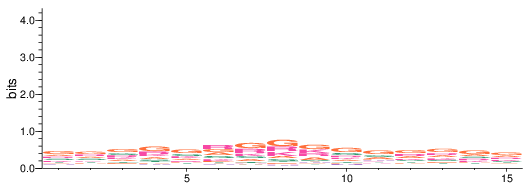

### filter_124.png

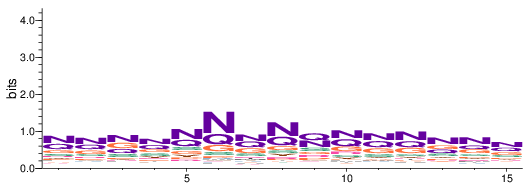

### filter_125.png

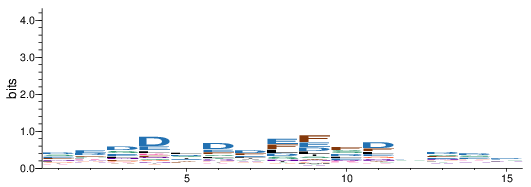

### filter_126.png

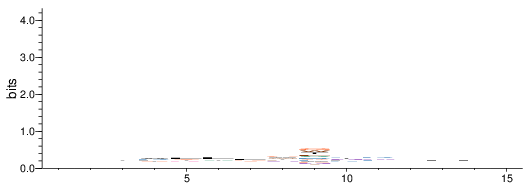

### filter_127.png

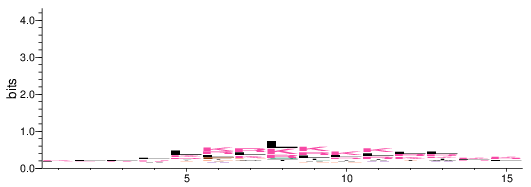

### filter_128.png

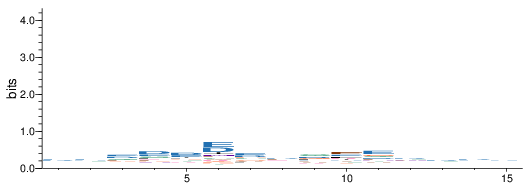

### filter_129.png

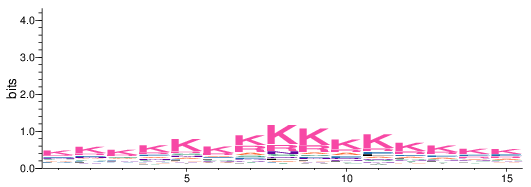

### filter_130.png

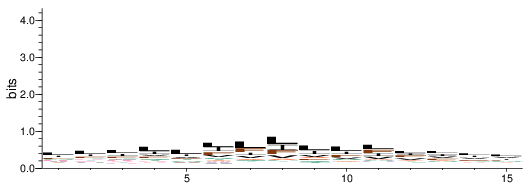

### filter_131.png

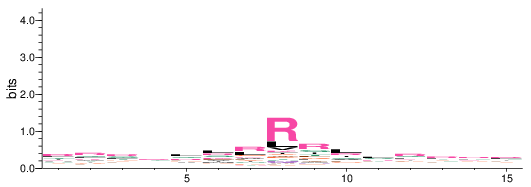

### filter_132.png

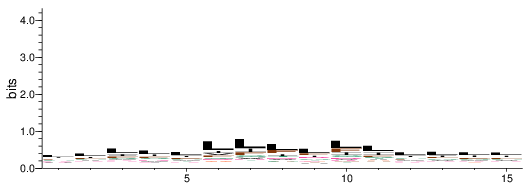

### filter_133.png

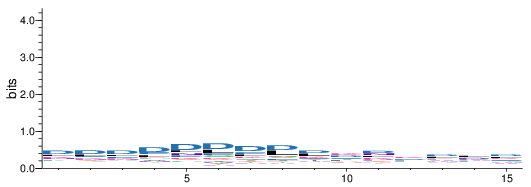

### filter_134.png

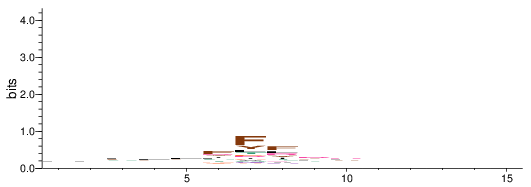

### filter_135.png

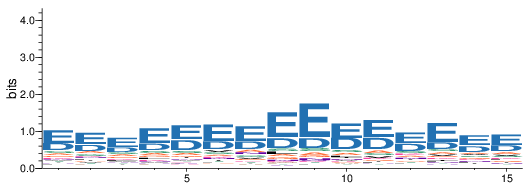

### filter_136.png

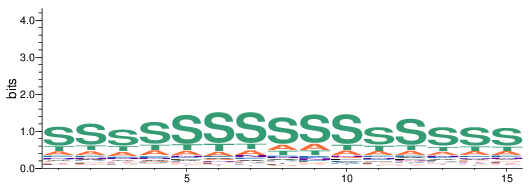

### filter_137.png

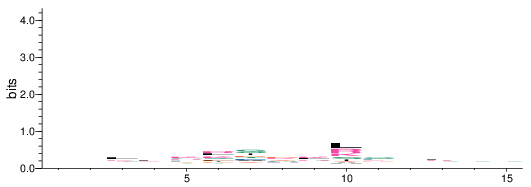

### filter_138.png

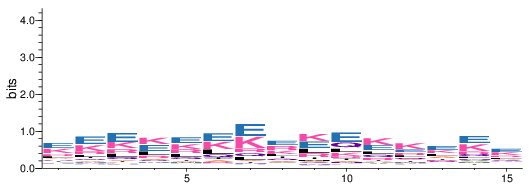

### filter_139.png

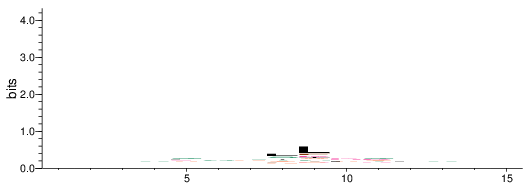

### filter_140.png

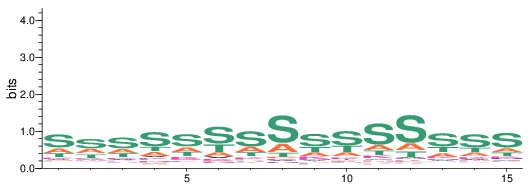

### filter_141.png

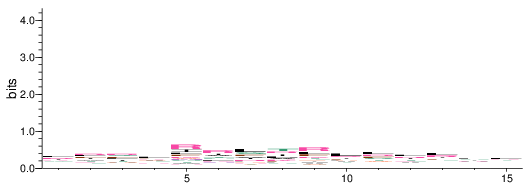

### filter_142.png

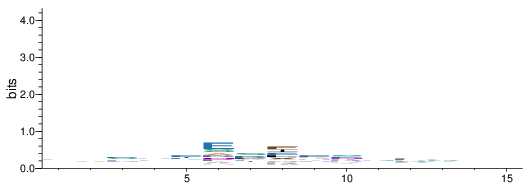

### filter_143.png

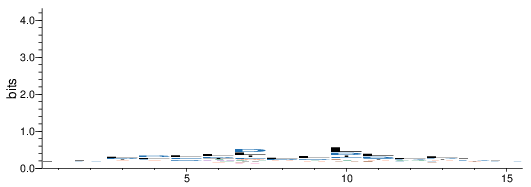

### filter_144.png

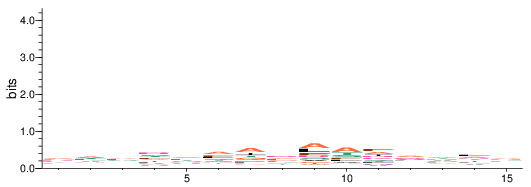

### filter_145.png

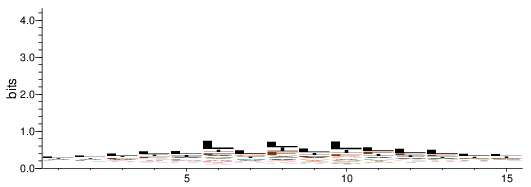

### filter_146.png

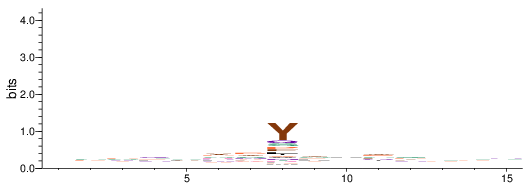

### filter_147.png

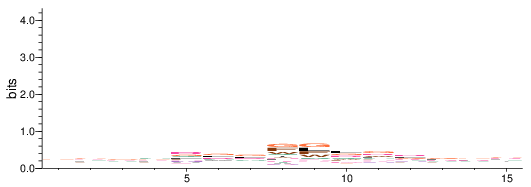

### filter_148.png

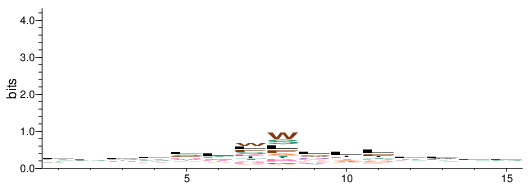

### filter_149.png

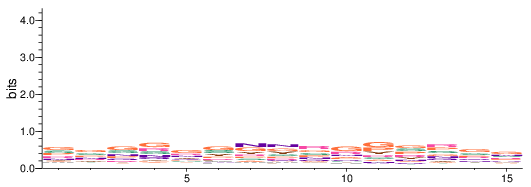

### filter_150.png

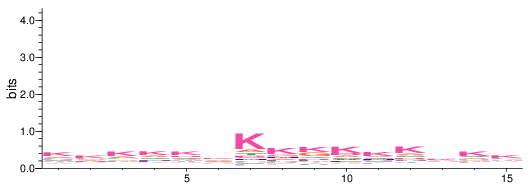

### filter_151.png

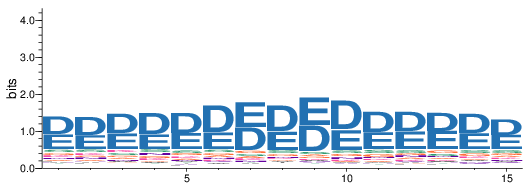

### filter_152.png

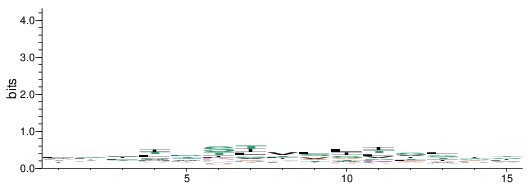

### filter_153.png

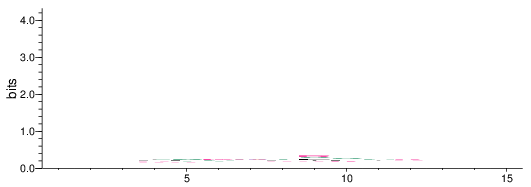

### filter_154.png

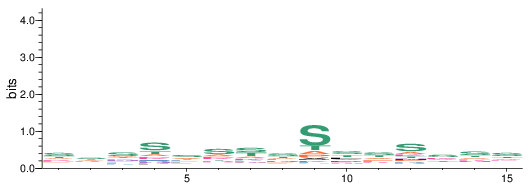

### filter_155.png

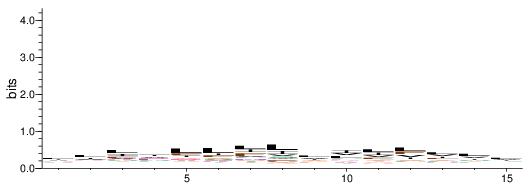

### filter_156.png

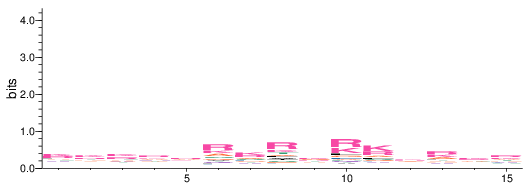

### filter_157.png

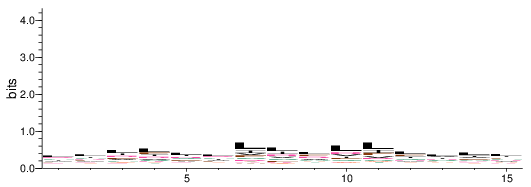

### filter_158.png

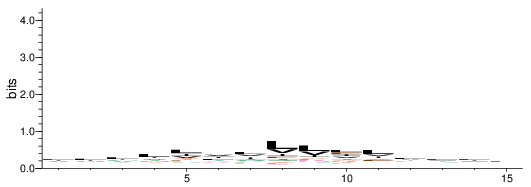

### filter_159.png

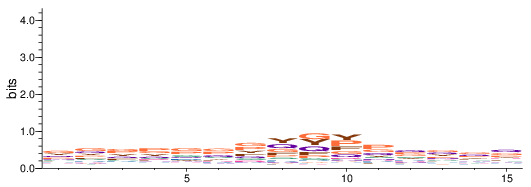

### filter_160.png

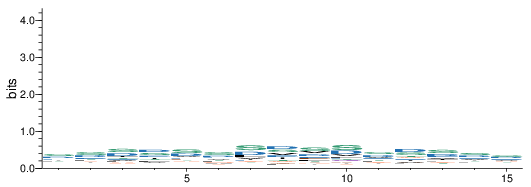

### filter_161.png

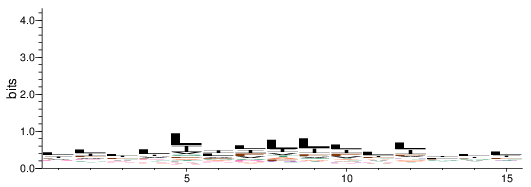

### filter_162.png

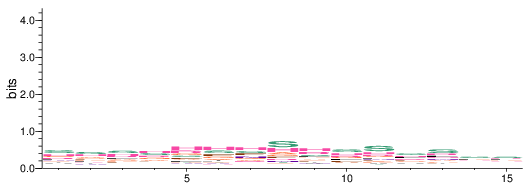

### filter_163.png

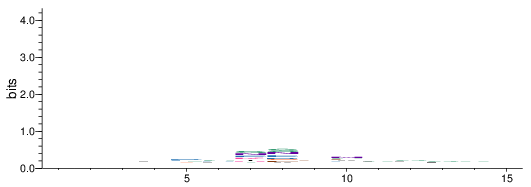

### filter_164.png

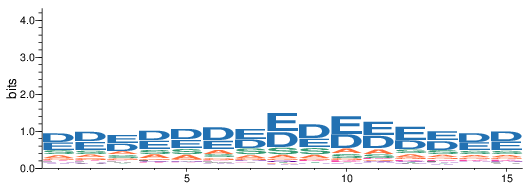

### filter_165.png

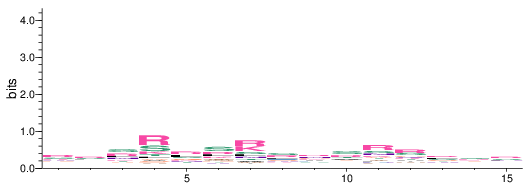

### filter_166.png

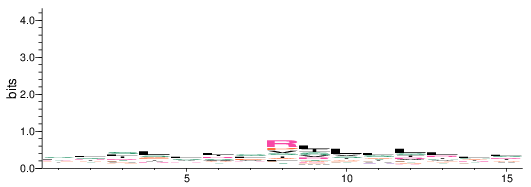

### filter_167.png

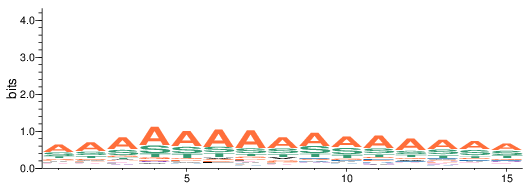

### filter_168.png

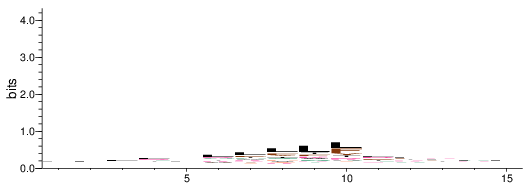

### filter_169.png

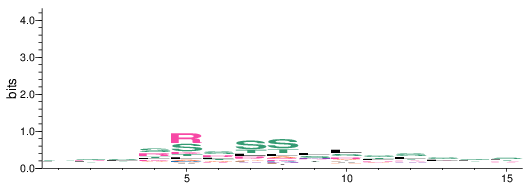

### filter_170.png

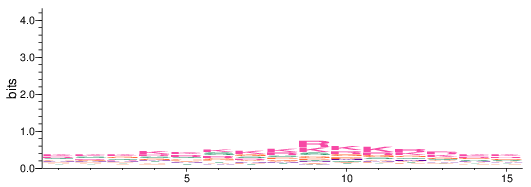

### filter_171.png

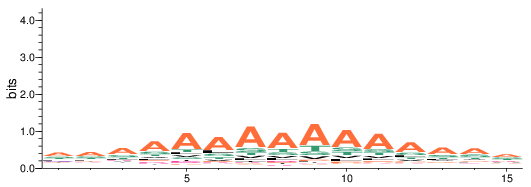

### filter_172.png

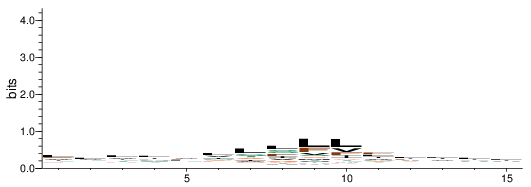

### filter_173.png

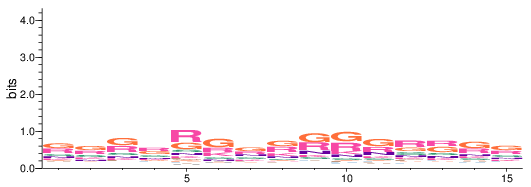

### filter_174.png

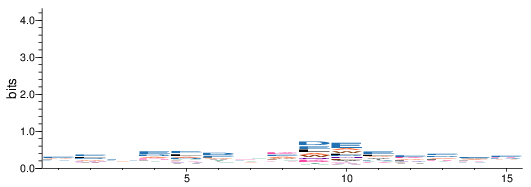

### filter_175.png

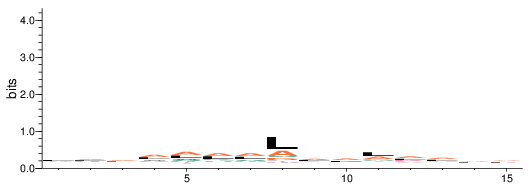

### filter_176.png

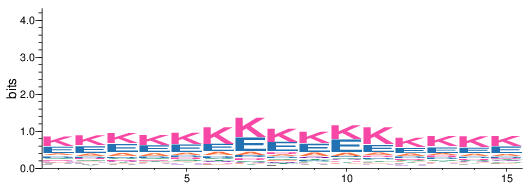

### filter_177.png

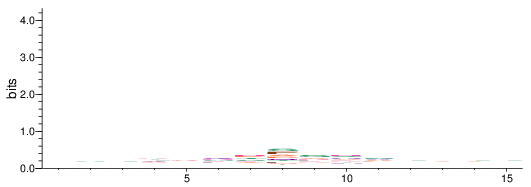

### filter_178.png

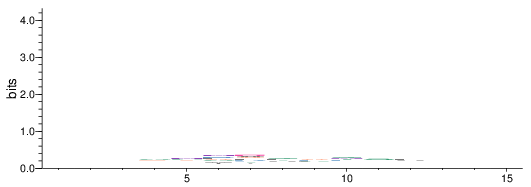

### filter_179.png

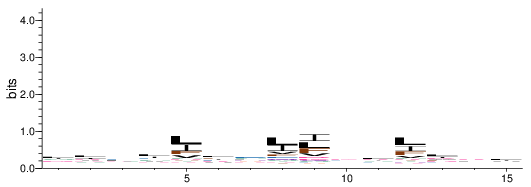

### filter_180.png

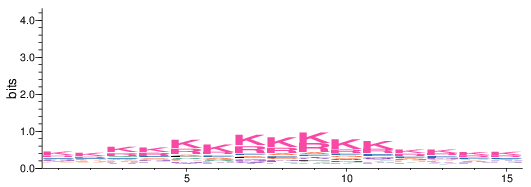

### filter_181.png

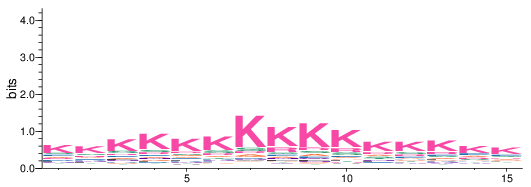

### filter_182.png

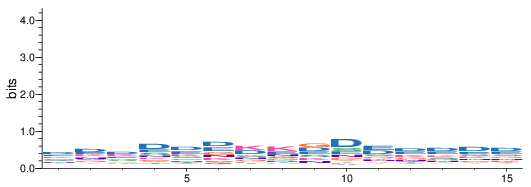

### filter_183.png

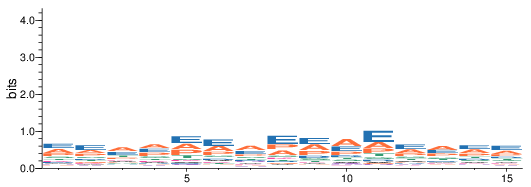

### filter_184.png

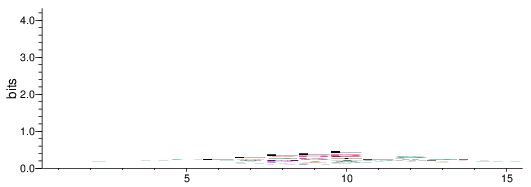

### filter_185.png

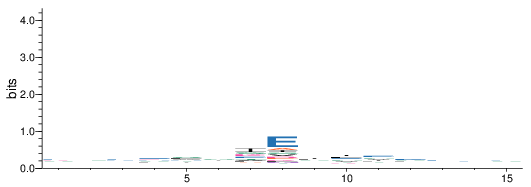

### filter_186.png

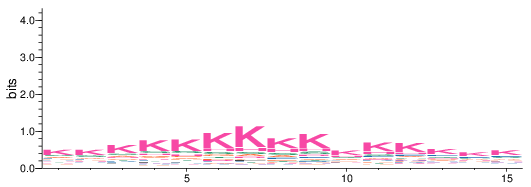

### filter_187.png

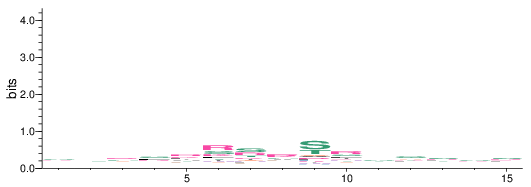

### filter_189.png

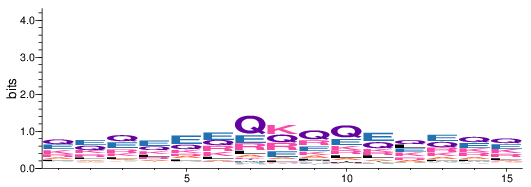
