## Supplementary Methods for "Discovering molecular features of intrinsically disordered regions by using evolution for contrastive learning"

**Reverse homology maximizes a lower bound on mutual information between sequences and their conserved functions in theory**

Our method joins a growing body of self-supervised work that uses the InfoNCE loss (and other related losses) for contrastive learning. In addition to its empirical performance in learning representations (Bachman, Hjelm, & Buchwalter, 2019; Chen, Kornblith, Norouzi, & Hinton, 2020; Hénaff et al., 2019; Hjelm et al., 2018; Oord, Li, & Vinyals, 2018), these losses lower-bound a maximization of the Shannon mutual information between input data and the output representation (Poole, Ozair, Oord, Alemi, & Tucker, 2019), i.e. $\text{max}_{g\in G}I(X,g\left( X \right))$ where $X$ is a random variable and $G$ is a set of deterministic functions (in these works, neural network encoders). These theoretical properties provide an explanation for why contrastive methods succeed at extracting relevant information from data. In this section, we will review some of these theoretical arguments, and use them to argue that reverse homology learns biologically sensible features.

Many contrastive methods apply the InfoNCE loss between different views of data: given input $x$, and transformations $t_{1}$ and $t_{2},$define ${v_{1}=t}_{1}(x)$ and ${v_{2}=t}_{2}(x)$ as two views. These transformations can be parameterless augmentations (e.g. in the context of images, rotations of the same image or with different illumination or contrast settings (Chen et al., 2020)), or another neural network summarizing the context of an image crop or portion of audio (Hjelm et al., 2018; Oord et al., 2018). Our method can be formulated similarly: in our case, $S_{q}$ is $v_{1}$and $s$ is $v_{2}$(we will follow the more general $v_{1}, v_{2}$notation in this section alone). In our work the homologous sequences we use to construct our query and target sets can be considered "evolutionary augmented" views of a common ancestor sequence $x$.

The intuition is that there is shared latent information between views that will be learned by the model: for example, when using augmentations to produce views, the objects in the image remain the same, but are just shown at different brightness or angles (Chen et al., 2020). Formalizing this intuition, van den Oord *et al*. show that optimizing the InfoNCE loss maximizes a lower bound on the mutual information between encoded views, $I(g_{1}\left( v_{1} \right);g_{2}\left( v_{2} \right))$ (Oord et al., 2018), suggesting that inputs with shared latent information (e.g. when two images contain the same object) will be correlated in the representation space. Second, Poole *et al*. show that maximizing this lower bound on mutual information between encoded views also means maximizing a lower bound on the mutual information between an encoded view and the original data $I(g\left( V \right);V)$ (Poole et al., 2019), suggesting that the encoders will learn to extract a representation from data that optimizes this correlation.

Tian *et al*. further formalize the "InfoMin" principle for selecting optimal views for contrastive learning (Tian et al., 2020). Ideally, we would like the model to learn relevant signal (like concepts about objects in images), while throwing away noise that is not informative for our analyses and applications (like incidental illumination or the viewing angle of the object.) The authors theoretically and empirically demonstrate that this kind of learning depends on training the model on good sets of view that share only the minimal necessary mutual information: views should share information we care about for downstream tasks, but should not be correlated in nuisance information.

Applying these ideas to sequences, a good set of views is two sequences that are highly diverged at the sequence level, but still carry out the same biological function (or otherwise similar biologically). We argue that using evolutionary homologues is a way to curate these kinds of views, especially when working with IDR homologues due to their rapid rate of evolution while preserving key properties essential to function (Zarin et al., 2019). When trained with this data, our model is expected to learn encoders that extract features that maximize the correlation between sequences with evolutionarily conserved properties.

**Mutational scanning sequence logos**

To visualize the mutational scanning maps as sequence logos, we first sum up the values for the change in feature for mutating the residue to each other amino acid at each position. This yields a negative number if mutating the residue is generally disfavored (i.e. most other residues at that position cause a drop in the feature), and a positive number if mutating the residue is generally favored (i.e. most other residues at that position cause an increase in the feature.) We use this sum as the scale for the total size of the letters in the sequence logo: we visualize negative positions as above the axis and positive positions as below the axis. In other words, the size of the letters correspond to the total magnitude of the drop or increase caused by mutating that amino acid to all other amino acids.

To visualize which specific amino acids are favored or disfavored at a specific position, we convert the vector of amino acids to a probability distribution. Since each individual feature has its own scale and average and max features especially exist on different scales, we first standardize the vectors by rescaling them between 0 and 10 with linear interpolation. For positions with a negative magnitude, we want to show the most favored amino acids (i.e. the amino acids that would induce the least drop in the feature, including synonymous mutations), so we calculate the Softmax over all amino acids at each position:

$$p\left( {AA}_{i} \right)=\frac{exp({AA}_{i})}{\sum_{j=1}^{20} {exp(AA}_{j})}$$

For positions with a positive magnitude, we want to show the least favored amino acids (i.e. the amino acids that would induce the highest increase in the feature), so we calculate the Softmax over all amino acids at each position but with negative terms:

$$p\left( {AA}_{i} \right)=\frac{exp({-AA}_{i})}{\sum_{j=1}^{20} {exp(-AA}_{j})}$$

Amino acids are then assigned a proportion of the magnitude in the visualization according to their probability with these formulas.
