## Supplementary figures and images for "Discovering molecular features of intrinsically disordered regions by using evolution for contrastive learning"

### filter_188.png

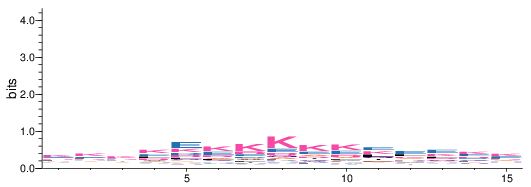

### filter_189.png

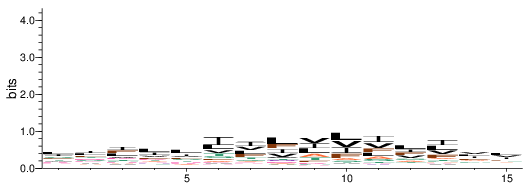
